# MK2a inhibitor CMPD1 abrogates chikungunya virus infection by modulating actin remodeling pathway

**DOI:** 10.1101/2021.05.26.445768

**Authors:** Prabhudutta Mamidi, Tapas Kumar Nayak, Abhishek Kumar, Sameer Kumar, Sanchari Chatterjee, Saikat De, Ankita Datey, Eshna Laha, Amrita Ray, Subhasis Chattopadhyay, Soma Chattopadhyay

## Abstract

Chikungunya virus (CHIKV) epidemics around the world have created public health concern with the unavailability of effective drugs and vaccines. This emphasizes the need for molecular understanding of host-virus interactions for developing effective targeted antivirals. Microarray analysis was carried out using CHIKV strain (Prototype and Indian) infected Vero cells and two host isozymes, MK2 and MK3 were selected for further analysis. Gene silencing and drug treatment were performed *in vitro* and *in vivo* to unravel the role of MK2/MK3 in CHIKV infection. Gene silencing of MK2 and MK3 abrogated around 58% CHIKV progeny release from the host cell and a MK2 activation (a) inhibitor (CMPD1) treatment demonstrated 68% inhibition of viral infection suggesting a major role of MAPKAPKs during the late phase of CHIKV infection *in vitro*. Further, it was observed that the inhibition in viral infection is primarily due to the abrogation of lamellipodium formation through modulation of factors involved in the actin cytoskeleton remodeling pathway that is responsible for releasing the virus from the infected cells. Moreover, CHIKV-infected C57BL/6 mice demonstrated reduction in the viral copy number, lessened disease score and better survivability after CMPD1 treatment. In addition, reduction in expression of key pro-inflammatory mediators such as CXCL13, RAGE, FGF, MMP9 and increase in HGF (a CHIKV infection recovery marker) was observed indicating the effectiveness of this drug against CHIKV. Additionally, CMPD1 also inhibited HSV1 and SARS CoV2-19 infection *in vitro*. Taken together it can be proposed that MK2 and MK3 are crucial host factors for CHIKV infection and can be considered as key targets for developing effective anti-CHIKV strategies in future.

**Author summary:** Chikungunya virus has been a dreaded disease from the first time it occurred in 1952 Tanzania. Since then it has been affecting the different parts of the world at different time periods in large scale. It is typically transmitted to humans by bites of *Aedes aegypti* and *Aedes albopictus* mosquitoes. Although, studies have been undertaken to combat the disease still there are no effective strategies like vaccines or antivirals against it. Therefore it is essential to understand the virus and host interaction to overcome this hurdle. In this study two host factors MK2 and MK3 have been taken into consideration to see how they regulate the multiplication of the virus. The *in vitro* experiments demonstrated that inhibition of MK2 and MK3 restricted viral infection Further, it was observed that this is due to the blocking of lamellipodium formation by modifying the factors involved in the actin cytoskeleton remodeling pathway that is responsible for releasing the virus from the infected cells. Besides, decreased disease score as well as better survivability was noticed in the *in vivo* experiments with mice. Therefore, MK2 and MK3 could be considered as the key targets for controlling CHIKV infection.

## Introduction

The Chikungunya virus (CHIKV) is an insect-borne virus belonging to the genus *Alphavirus* and family *Togaviridae* and transmitted to humans by *Aedes* mosquitoes(1). Three CHIKV genotypes, namely West African, East Central South African and Asian have been identified. The incubation period ranges from two to five days following which symptoms such as fever (up to 40°C), petechial or maculopapular rash of the trunk and arthralgia affecting multiple joints develop(2–4).

CHIKV is a small (60-70nm diameter), spherical, enveloped, positive sense single-stranded RNA (~12Kb) virus (5–7). Its genomic organization is 5’-cap-nsP1-nsP2-nsP3-nsP4-(junction region)-C-E3-E2-6K-E1-3’(8). The non-structural proteins (nsP1-4) are primarily involved in virus replication, while structural proteins C, E3, E2, 6K and E1 are responsible for packaging and producing new virions.

In India, CHIKV infection has re-emerged with the outbreak of 2005–08 affecting approximately 1.3 million people in 13 different states (9). The clinical manifestations during these outbreaks were found to be more severe leading to the speculation that either a more virulent or an efficiently transmitted variant of this virus might have emerged (10).

CHIKV, among most other viruses across families, interacts with a number of cellular proteins and consequently metabolic pathways to aid its survival in the host (11–17). Several facets of CHIKV pertaining to strategies required for ecological success, replication, host interaction and genetic evolution are yet to be fully explored and are constantly evolving. This spurs the need to identify important host pathways that can be targeted for developing antiviral therapies against the virus.

Alternatively, host factors involved in viral replication may also be targeted. Previous studies have shown compounds targeting furin, protein kinases, and Hsp90, are inhibiting CHIKV replication *in vitro* (18–20). However, further validation through *in vivo* experiments and pre-clinical studies need to be performed prior to developing effective antivirals.

Our group has formerly reported an Indian outbreak strain IS, to exhibit a faster replication rate than the CHIKV prototype strain, PS *in vitro* (21). The present study identifies host genes which are modulated differentially during CHIKV infection in mammalian system and explores the involvement of MAPK-activated protein kinases during virus infection using both *in vitro* and *in vivo* conditions through inhibitor studies.

## Materials and Methods

### Cells, Viruses, Antibodies, Inhibitors

Vero cells (African green monkey kidney cells), CHIKV strains, prototype strain, PS **(Accession no: AF369024.2)** and novel Indian ECSA strain, IS **(Accession no: EF210157.2)** and E2 Monoclonal antibody were gifted by Dr. M. M. Parida, DRDE, Gwalior, India. The HSV-1 virus strain KOS with GenBank accession Number JQ673480.1 was kindly gifted by Dr. Roger Everett, Glasgow University, Scotland. The HEK 293T cell line was gifted by Dr. Rupesh Dash, Institute of Life Sciences, Bhubaneswar, India. **SARS details: -** The SARS-CoV-2 virus used in this study was isolated from a clinically confirmed local COVID-19 patient (GISAID accession ID-EPI_ISL_1196305). Virus from the 10^th^ passage was used for experiments. Cells were maintained in Dulbecco’s Modified Eagle’s medium (DMEM; PAN Biotech, Germany) supplemented with 5% Fetal Bovine Serum (FBS; PAN Biotech), Gentamicin and Penicillin-Streptomycin (Sigma, USA). The anti-nsP2 monoclonal antibody used in the experiments was developed by us(22). Cofilin monoclonal antibody was purchased from Cell Signaling Technologies (Cell Signaling Inc, USA). The pMK2 polyclonal antibody and MK3 monoclonal antibody were purchased from Santacruz Biotechnology (USA). The p-Cofilin antibody and GAPDH antibody were procured from Sigma Aldrich (USA) and Abgenex India Pvt. Ltd. (India) respectively. Anti-mouse and anti-rabbit HRP-conjugated secondary antibodies were purchased from Promega (USA). Alexa Fluor 488 and Alexa Fluor 594 antibodies were purchased from Invitrogen (USA). The MK2a inhibitor, CMPD1 was purchased from Calbiochem (Germany).

### Virus infection

The Vero cells were infected with PS/IS strains of CHIKV respectively according to the experimental requirements as reported earlier (21). Thereafter, CHIKV infected cells were incubated for 15-18hpi following which cells and supernatants were harvested from mock, infected and drug treated samples for downstream processing. For HSV (Herpes Simplex Virus) and SARS-CoV-2, Vero cells were infected with 0.1 MOI of virus and incubated for 22 hpi. The supernatants were harvested at 22 hpi and subsequent downstream processing was carried out for estimating the viral titers.

### RNA isolation and Microarray hybridization

In the present study, the global gene expression analyses were carried out using Agilent Rhesus GeneChip® ST arrays. Sample preparation was performed according to the manufacturer’s instruction (Agilent, USA). Briefly, RNA was extracted from mock and virus infected Vero cells using the RNeasy mini kit (Qiagen, Germany). Next, RNA quality was assessed by Agilent Bioanalyzer and cDNA was prepared using oligo dT primer incorporating a T7 promoter. The amplified, biotinylated and fragmented sense-strand DNA targets were generated from the extracted RNA and hybridized to the gene chip containing over 22,500 probe sets at 65°C for 17h at 10 rpm. After hybridization, the chips were stained, washed and scanned using a Gene Chip Array scanner.(23)

### Microarray analysis

Raw data sets were extracted from all text files after scanning the TIFF files. These raw data sets were analyzed separately using the GeneSpring GX12.0 software (Agilent Technologies, USA) followed by differential gene expression and cluster analysis. Differential gene expression analyses were performed by using standard fold change cut off >=2.0 and >=10.0 against IS (8hpi) vs Mock (8hpi), PS (8hpi) Vs Mock (8hpi), PS (18hpi) Vs Mock (18hpi), IS (8hpi) vs PS (8hpi) and IS (8hpi) vs PS (18hpi). The hierarchical clustering was performed using the Genesis software(24). Functional annotation of differentially expressed genes was carried out using the PANTHER gene ontology analysis software (25).

### RNA extraction and qRT-PCR

Equal volumes of serum isolated from all groups of mice samples were taken for viral RNA isolation using the QiaAmpViral RNA isolation kit (Qiagen, USA) as per the manufacturer’s instructions. RT reaction was performed with 1 μg RNA using the First Strand cDNA Synthesis kit (Fermentas, USA) as per manufacturer’s instructions. Equal volume of cDNA was used for PCR amplification of E1 gene of CHIKV using specific primers (26). The nucleocapsid (NC) gene of SARS-CoV-2 was amplified using forward primer-5’-GTAACACAAGCTTTCGGCAG-3’ and reverse primer-5’-GTGTGACTTCCATGCCAATG-3’. The viral copy number estimation from Ct values was estimated from the standard curve generated for CHIKV E1 gene/ SARS-CoV-2 (NC) gene (data not shown).

### siRNA Transfection

Monolayers of HEK 293T cells with 70% confluency (1×10^6^ cells/well) in 6-well plates were transfected separately or in combination with 60pmols of siRNA corresponding to MK2 mRNA sequence [(5’-3’)CCAUCACCGAGUUUAUGAAdTdT] and MK3 mRNA sequence [<5’-3’> GAGAAGCUGCAGAGAUAAUdTdT] or with siRNA negative control. Transfection was performed using Lipofectamine-2000 (Invitrogen, USA) according to the manufacturer instructions. In brief, HEK cells were transfected using Lipofectamine 2000 according to different siRNA quantity in Opti-MEM medium (Thermo scientific, USA). The transfected cells were infected with either CHIKV strains PS or IS with MOI 0.1 at 24 hours post transfection (hpt). Eighteen hours post infection, the cells were harvested to measure the nsP2 and MK2/3 protein levels by Western blot analysis.

### SDS-PAGE and Western blot analysis

Protein expression was examined by Western blot analysis as described earlier (21, 27). CHIKV nsP2 and E2 proteins were detected with monoclonal antibodies (28) and re-probed with GAPDH antibody to confirm the equal loading of samples. The pMK2, MK3, Cofilin and pCofilin antibodies were used as recommended by the manufacturer. The Western blots were scanned using the Quantity One Software (Bio Rad, USA).

### Plaque assay

The CHIKV-infected cell culture supernatants were collected at 18 hpi and subjected to plaque assay according to the procedure mentioned earlier (29).

### Immunofluorescence staining

Immunofluorescence staining was carried out using the procedures described earlier (22). Vero cells were grown on glass coverslips placed in 35mm dishes and infected with CHIKV (MOI 0.1) as described above. At 18 hpi, coverslips were stained with primary antibodies followed by staining with secondary antibody (AF 594-conjugated anti-mouse antibody) for 45 mins. The phalloidin staining was carried out using the Cytopainter F actin labeling kit as per manufacturer’s protocol (Abcam, UK). The coverslips were stained with DAPI for 90 sec and mounted with 15-20 μl Antifade (Invitrogen, USA) to reduce photo-bleaching. Fluorescence microscopic images were acquired using the Leica TCS SP5 confocal microscope (Leica Microsystems, Germany) with 63X objective and analyzed using the Leica Application Suite Advanced Fluorescence (LASAF) V.1.8.1 software.

### Immunohistochemistry analysis

For histopathological examinations, tissue samples were dehydrated, embedded in paraffin wax, and thereafter serial paraffin sections (5μm) were obtained (30). Briefly, the sections were immersed in two consecutive xylene washes for de-paraffinization and were subsequently hydrated with five consecutive ethanol washes in descending order of concentration: 100%, 90%, 70%, and deionized water. The paraffin sections were then stained with hematoxylin-eosin (H&E), and histopathological changes were visualized using a light microscope (Zeiss Vert.A1, Germany).

### Cellular cytotoxicity assay

Cellular cytotoxicity assay was performed as described earlier (31). Vero cells were seeded onto 96-well plates at a density of 3000 cells/well, treated with different concentrations of CMPD1 for 24 hrs at 37°C with 5% CO_2_. DMSO-treated samples served as control. After incubation, 10μl of MTT reagent (Sigma Aldrich, USA) was added to the wells followed by incubation at 37°C for 3hrs and processed further. Absorbance of the suspension was measured at 570nm using ELISA plate reader (BioRad, USA). Cellular cytotoxicity was determined in duplicates and each experiment was repeated thrice independently.

### CMPD1 treatment

Vero cells with 90% confluency were grown in 35mm or 60mm cell culture dishes (according to the experimental requirements) and infected with PS or IS strains of CHIKV as described above at MOI 0.1. After infection, cells were treated with either DMSO or different concentrations of CMPD1 as per the protocol mentioned earlier (32). The cells were observed for detection of cytopathic effect (CPE) under 10X objective of bright field microscope. Infected cells and supernatants were then collected at 15-18hpi depending on the experiment.

### Time of addition experiment

Vero cells were infected with CHIKV as described above and CMPD1 (50μM) was added at 1hr interval upto 11hrs to the infected cells in different dishes. Thereafter, cell culture supernatants of all the samples were harvested at 15hpi and plaque assay was carried out for estimating viral titer.

### CHIKV infection in mice

The mice related experiments were performed as per CPCSEA guidelines and were approved by the IAEC committee. Around 10-14 days old male C57BL/6 mice (n=5) were injected subcutaneously with 1×10^6^ particles of IS in DMEM. At 3hpi, mice were fed with CMPD1 at a concentration of 5mg/kg of body weight and continually fed at every 24hr-interval up to3 days. All mice were sacrificed on the fourth day; blood samples were harvested from mock, infected and drug-treated samples and used for downstream processing. For survival curve analysis, CHIKV-infected mice were fed with CMPD1 and observed every day, for CHIKV-induced disease manifestations up to 8 days post infection (dpi). All infected mice were scored on a scale of 0 to 6 based on CHIKV induced disease symptoms such as(0- No symptoms, 1- lethargic, 2- ruffled fur, 3- restricted movement/limping, 4- one hind limb paralysis and 5 – both hind limb paralysis 6- Morbid/dead).

### Proteome profiling

In order to assess the levels of different cytokines in mock, CHIKV-infected and CHIKV-infected+drug treated mice samples, proteome profiling was performed using the Mouse XL cytokine array kit (R & D systems, USA) as per manufacturer’s instructions. The array blots were incubated with serum samples at 4°C overnight on a gel rocker, followed by incubation with HRP-conjugated secondary antibody. Blots were developed using the chemiluminescent HRP substrate and scanned by the Image Lab software (Bio-Rad, USA). The relative differences in expression patterns of selected cytokines among the different groups of samples were assessed using the GraphPadPrism8 software.

#### Bioavailability Prediction

The bioavailability of CMPD1 was predicted through the SWISS ADME tool available in the website (www.swissadme.ch). The SMILE structure of CMPD1 was submitted to the tool for analysis and prediction.

### Statistical analysis

Statistical analysis of the experimental data was performed by using the GraphPad Prism 8.0 Software and presented as mean±SD of three independent experiments. The One-way ANOVA with Dunnet post-hoc test was used to compare the differences between the groups. In all the tests, *p* value < 0.05 was considered to be statistically significant.

### Accession numbers

The accession number for the submitted microarray experimental data to Array Express database is **E-MTAB-6645.** The URL for the submitted microarray experimental data is as follows: *http://www.ebi.ac.uk/arrayexpress/experiments/E-MTAB-6645.*

## Results

### Differential host gene expressions for PS and IS strains of CHIKV in Vero cells

Earlier, it was observed that CPE developed by IS was more prominent at 8 hpi as compared to PS which showed similar CPE around 18 hpi(21). To understand the host gene expression profiles for the two CHIKV strains, Vero cells were harvested at 8 and 18 hpi for microarray analysis. Microarray data revealed the differential expression of 20227 genes, of which 12221 genes were differentially expressed after applying fold change cut off ≥2.0. Further, 684 genes from the 12221 were differentially expressed with fold change ≥10.0. The cluster analysis of differentially expressed genes was carried out using the GENESPRING GX 12.0 software, as shown in Fig 1A. Annotation of the total genes into different protein classes was carried out using the Panther software. It was observed that majority of the genes belonged to the nucleic acid binding molecules, signaling molecules, transcription factors among others, as represented in Fig 1B. A pie-chart was constructed using the Panther software to annotate these genes into different biological processes, and it was observed that majority of the modulated genes belonged to the pathways involved in different cellular processes (Fig 1C). Moreover, 720 genes were differently regulated by IS alone as depicted by the Venn diagram constructed through the Gene Venn software in (Fig 1D and 1E). Out of these 720 genes, few selected genes were functionally annotated into different host cellular pathways as shown in “S1 Table”. MK3 was present among the 720 genes that were antagonically expressed in IS infected cells at 8 hpi in comparison to PS (8 and 18 hpi). The importance of MK3 and its isozyme partner MK2 was thus, deliberated during CHIKV infection in this study. Together, the data indicate that CHIKV utilizes different host cell pathways for efficient replication inside the host cell and there are differential host gene expression patterns for various strains of CHIKV.

**Fig 1:**
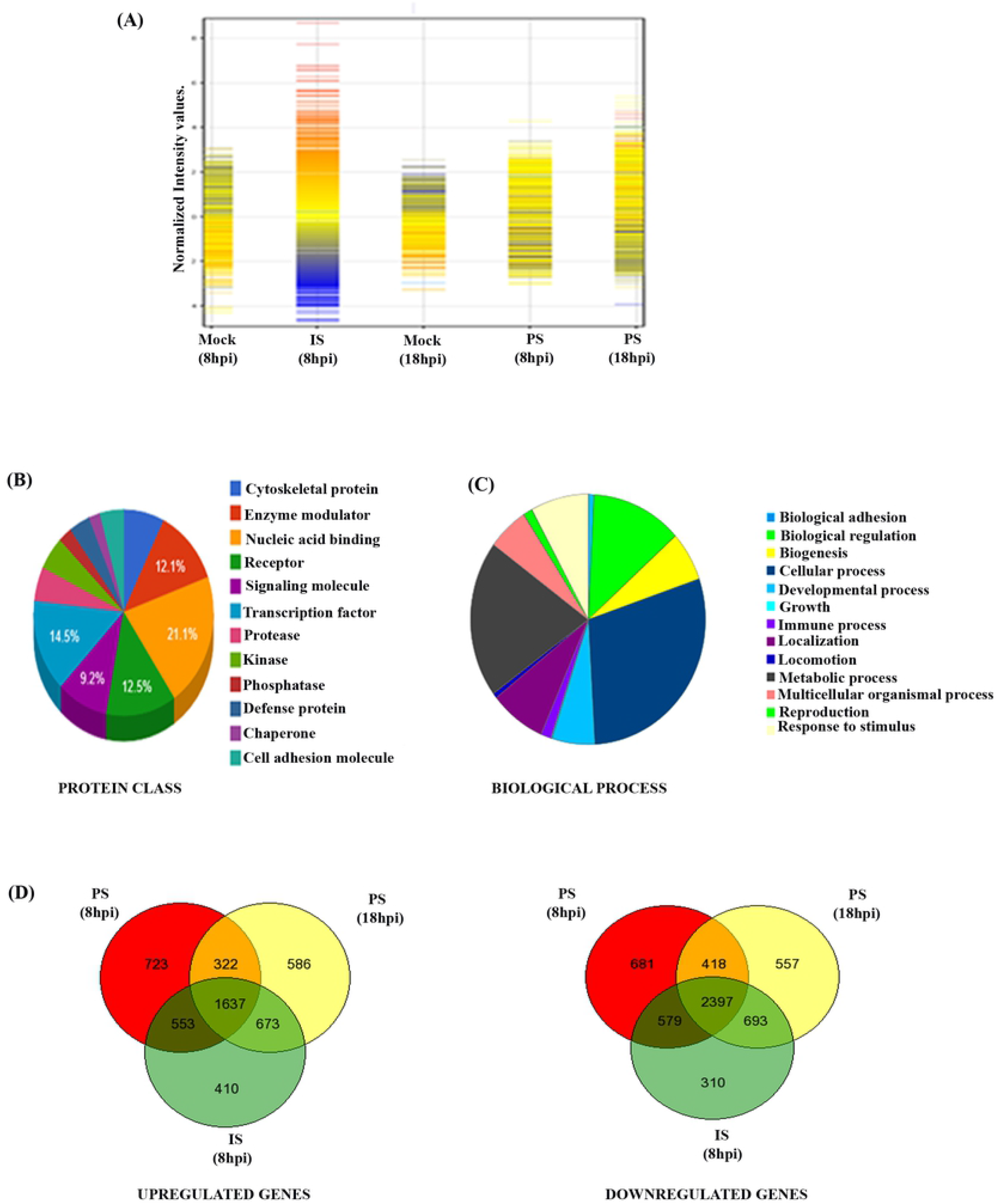
Differential host gene expressions for PS and IS strains of CHIKV in Vero cells. **(A)** Hierarchical clustering showing the overall expression patterns of the modulated host genes by PS/IS strains of CHIKV during infection in mammalian cells. **(B)** Pie-chart depicting the distribution of the host genes in CHIKV-infected samples into different protein classes. **(C)** Pie-chart depicting the distribution of the modulated host genes into different cellular processes. **(D and E)** Venn diagram showing both commonly and differentially regulated host genes in CHIKV (PS/IS) infected Vero cells.

### MK2 and MK3 gene silencing abrogates CHIKV progeny release without affecting viral protein synthesis

To elucidate the importance of MK2 and MK3 in CHIKV infection, gene silencing through siRNA approach was employed. Since the transfection efficiency of Vero cells is poor, HEK293T cell line (Kidney epithelial cell line) was used for this experiment. HEK293T cells were transfected with 60 pmol of MK2 and/or MK3 siRNAs and incubated for 24 hrs at 37°C.

Next, the siRNA transfected cells were infected with CHIKV [(PS/IS), MOI 0.1] and cells as well as supernatants were harvested at 18 hpi for further analysis. No remarkable change in nsP2 expression was observed after genetic knock down of either MK2 or MK3. Surprisingly, the expression of CHIKV-nsP2 was increased marginally when both MK2 and MK3 were silenced together as compared to control as shown in Fig 2A and 2B (left and right panels respectively).

**Fig 2:**
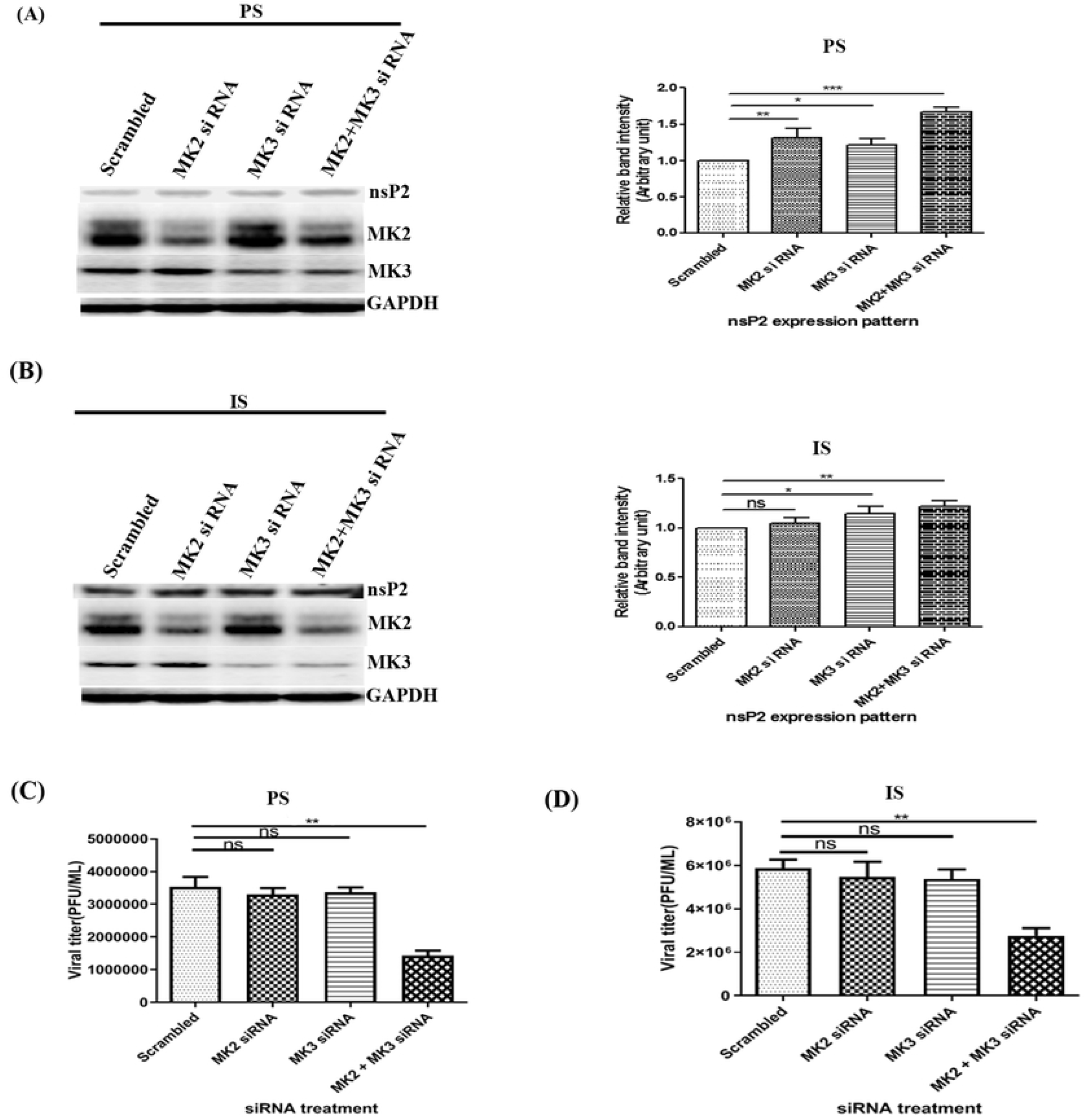
MK2 and MK3 gene silencing abrogates CHIKV progeny release without affecting viral protein synthesis. (**A and B)** After 24 hrs post transfection with 60 pmol of MK2/3 siRNA (either separately/in combination), cells were super-infected (PS/IS MOI 0.1) and harvested at 18 hpi. Western blot showing the expression levels of different proteins (Left panel). Bar diagrams showing relative band intensities of different proteins (Right panel). GAPDH was used as control. **(C and D)** Bar diagram showing the viral titres after siRNA treatment for PS and IS strains, (n=3; *p*<0.05).

As the expression of nsP2 was increased after siRNA treatment, plaque assay was performed to assess the effect of MK2 and/or MK3 down-regulation in viral progeny formation. Interestingly, it was observed that the viral titers were reduced by 58% for PS strain and 53% for IS strain as shown in Fig 2C and 2D. Therefore, it can be suggested that MK2 and MK3 altogether affects CHIKV progeny release without affecting viral protein synthesis.

### CMPD1, an MK2a inhibitor abrogates CHIKV infection *in vitro*

The MAPK-activated protein kinases MAPKAPK3 (MK3) and MAPKAPK2 (MK2) are the substrates of P38 MAPK that form a pair of structurally and functionally closely related enzymes. Being highly homologous enzymes (around 70% at the amino acid sequence), their substrate spectrums are indistinguishable(33). MK2 expression levels usually exceeds MK3 level in cells, however, in absence of functional MK2, MK3 compensates.

To investigate the role of MK2 pathway in CHIKV infection, Vero cells were treated with a non-ATP competitive MK2 inhibitor, CMPD1, which selectively inhibits P38-mediated MK2 activation (34). In order to determine the cytotoxicity of CMPD1, Vero cells were treated with different concentrations of the drug (25, 50, 75 and 100μM) for 24 h and MTT assay was performed. It was observed that 98%, 95% and 85% cells were viable with 25, 50 and 100μM concentrations of the drug, respectively, as shown in Figure 3a. Next, dose kinetics assay was performed to determine the anti-CHIKV efficacy of CMPD1. Therefore, Vero cells were infected with two different strains of CHIKV with MOI 0.1 and treated with 25, 50 and 100μM concentrations of CMPD1. The cell culture supernatants were harvested at 18 hpi and plaque assay was carried out to estimate the virus titers. Around 90% decrease in virus titer was observed with higher concentrations of CMPD1 in comparison to DMSO control for both the strains (Fig 3B and 3C). Since, effect of CMPD1 was same for both the strains, IS strain (more virulent of the two strains used in this study) was used for further experiments.

**Fig 3:**
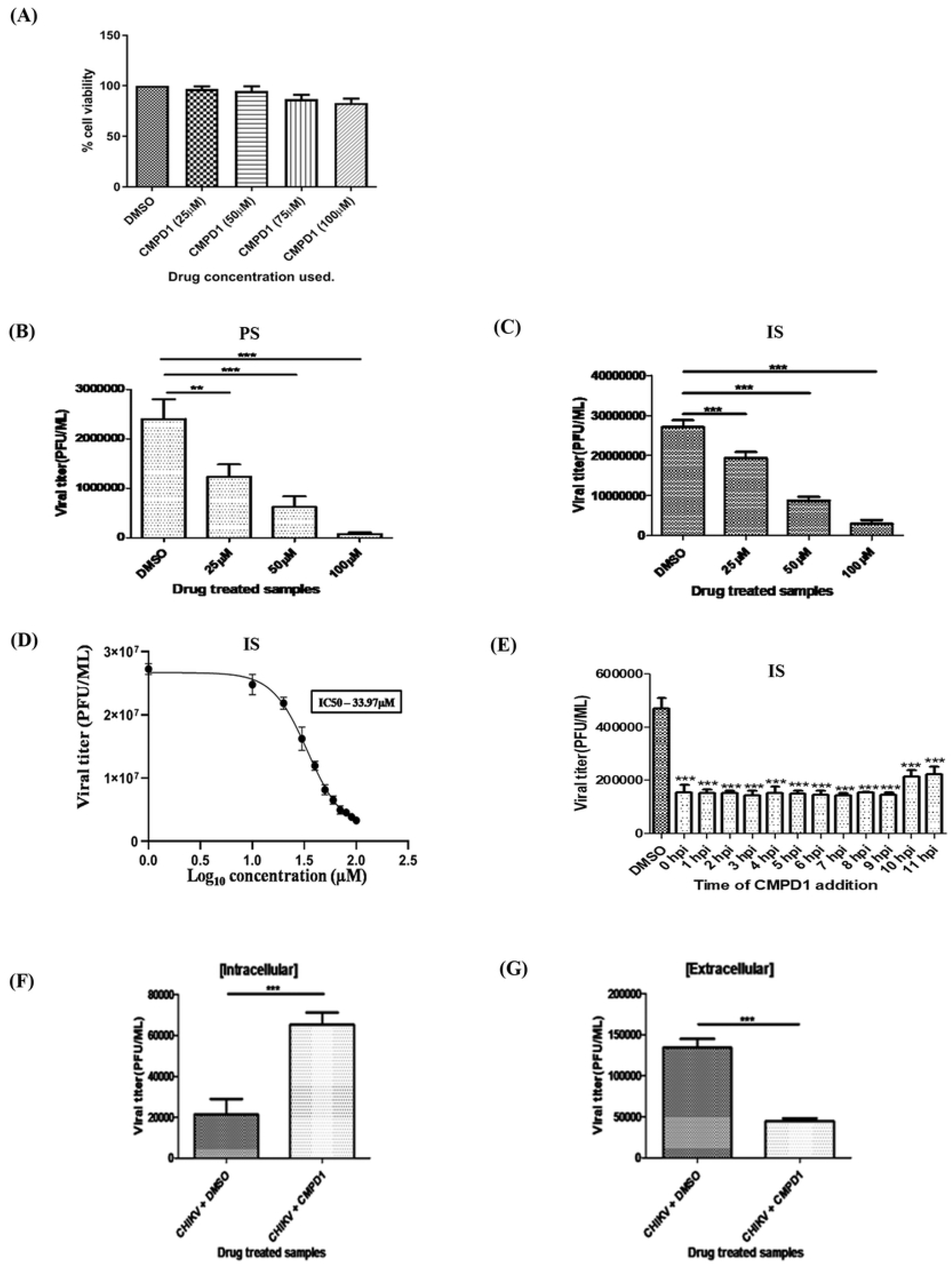
CMPD1, an MK2a inhibitor abrogates CHIKV infection *in vitro*. **(A)**Vero cells were treated with different concentrations of CMPD1 (25, 50, 75 and 100 μM) for 24 h and MTT assay was performed. **(B and C)** Vero cells infected with CHIKV PS/IS at MOI 0.1 and drug treated. Bar graph showing the viral titers in the presence of CMPD1 (25, 50 and 100 μM). **(D)** Dose response curve showing the IC50 of CMPD1 against CHIKV. **(E)** Bar graph showing the viral titers estimated through plaque assay from the supernatants obtained from the time of addition experiment for CMPD1(50μM) post CHIKV infection. **(F and G)** Bar graph showing intracellular and extracellular virus titers for samples harvested at 18hpi. DMSO was used as control. All the graphs depict the values of mean ± SD (\**p< 0.05*) of three independent experiments.

To estimate the IC_50_ value of CMPD1, Vero cells were infected with CHIKV as mentioned above and different concentrations of CMPD1 (10-100μM) were added to the cells post-infection. The supernatants were harvested at 18 hpi and plaque assay was performed. The plaque numbers were converted into log 10 of PFU/mL and plotted in the graph as shown in Fig 3D. The IC_50_ of CMPD1 was found to be 33.97 μM.

Next, to assess the possible mechanism of action of CMPD1 on CHIKV replication, time of addition experiment was performed. Vero cells were infected with IS strain with MOI 0.1 and 50 μM of CMPD1 was added at 1hr interval from 0–11 hpi. DMSO was used as a control. Next, the CHIKV-infected and CMPD1 treated supernatants were harvested at 15 hpi and plaque assay was performed as mentioned above to assess the release of infectious virus particles. As shown in Fig 3E, it was observed that around 55% of the infectious virus particle release was abrogated in the presence of 50 μM of CMPD1, even after the addition of the drug at 11 hpi. This indicates that CMPD1 inhibits later phase of CHIKV life cycle.

Lastly, to understand the role of CMPD1 in CHIKV packaging/release, Vero cells were infected, drug-treated, and supernatants were collected at 18 hpi for estimating the extracellular viral titer through plaque assay. For estimating the intracellular virus titer, cells were washed twice with 1X PBS and harvested. The pelleted cells were resuspended in fresh serum free medium and freeze-thawed thrice to release virus particles trapped inside the cells. Then the plaque assay was performed using the supernatant to estimate the intracellular virus titer. Similar to the previous experiment, the extracellular virus titer was around 70% less in CMPD1 treated samples in comparison to control but the intracellular virus titer was around 60% more for CMPD1 treated samples as shown in Fig 3F and 3G. This suggests that CMPD1 did not inhibit the formation/packaging of newly synthesized host particles inside the host cell; however, it affects the release of CHIKV viral progeny from the host cell.

### CMPD1 blocks the actin polymerization process modulated by CHIKV for its progeny release

It is well known that both the isozymes, MK2 and MK3 are exclusively phosphorylated by P38 MAPK and both have similar substrates (35). It is also known that LIM kinase 1 (LIMK1), a downstream substrate of MK2 induces actin polymerization by phosphorylating and inactivating cofilin, an actin-depolymerizing factor (36, 37). Therefore, to understand the effect of CMPD1 in viral infection and on downstream substrates of MK2, the cells were infected with IS at MOI 0.1 and treated with 50 μM CMPD1. Infected cells were observed for the development of CPE at 18 hpi and clear reduction in CPE was observed after CMPD1 treatment (Fig 4A). The cells were harvested at 18 hpi and cell lysates were processed for Western blot analysis. It was noticed that the levels of pMK2 and MK3 were downregulated after drug treatment with no change in CHIKV nsP2 expression as shown in Fig 4B and 4C. Similarly, the expression of Cofilin and p-Cofilin was decreased in the presence of CMPD1. The expression of pMK3 could not be tested due to unavailability of a commercial antibody. Altogether, the data suggest that MK2 phosphorylation plays an important role in viral progeny release by modulating the actin polymerization process.

**Fig 4:**
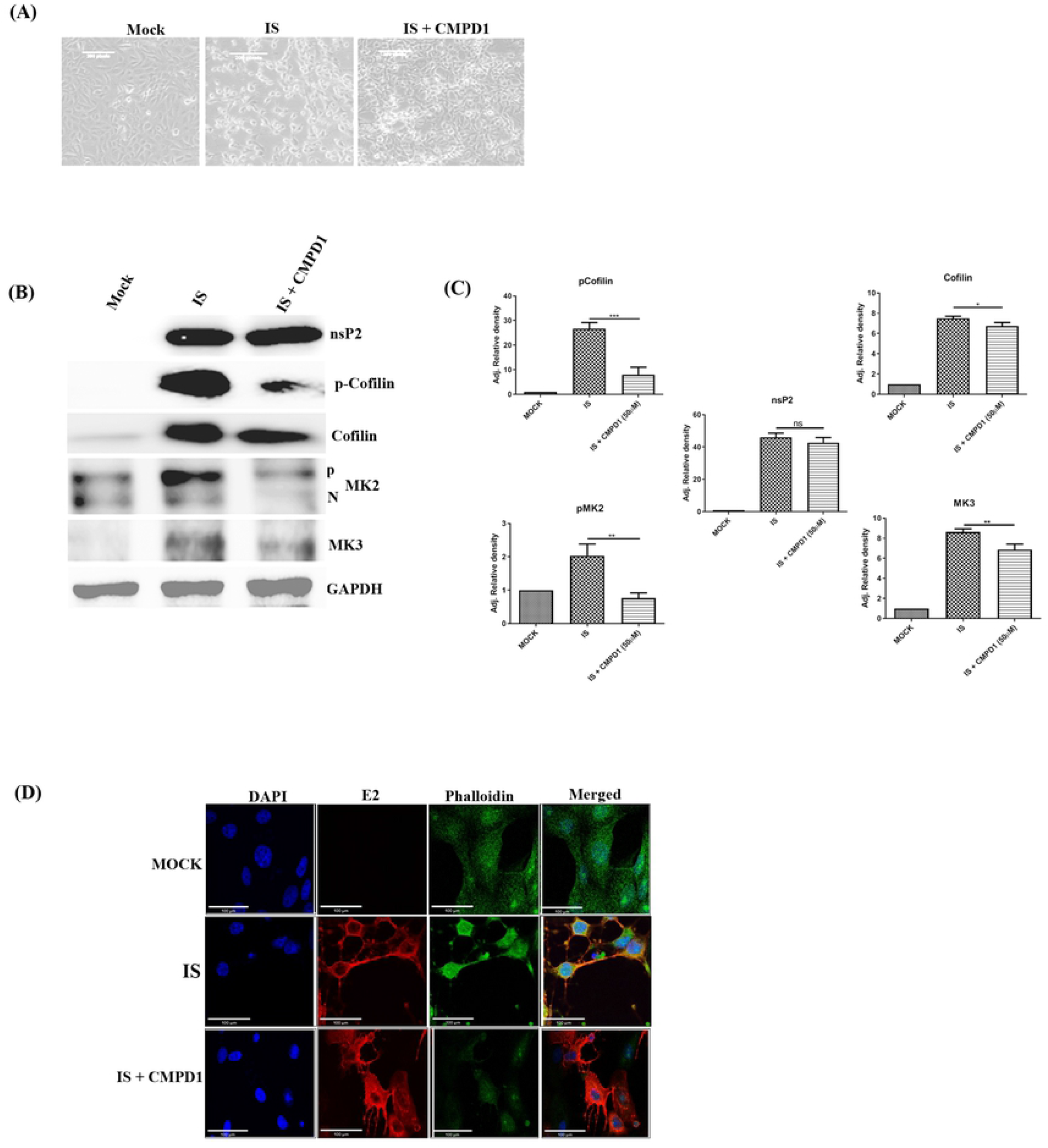
CMPD1 blocks the actin polymerization process modulated by CHIKV for its progeny release. Vero cells were infected with the IS strain (0.1 MOI), 50 μM of CMPD1 was added to the cells and incubated for 18 hpi. **(A)** Bright field images (20X magnification) showing the cytopathic effect after CHIKV infection with or without CMPD1 treatment (50 μM). **(B)** Western blot analysis showing the expressions of nsP2, pMK2, MK3, Cofilin and p-Cofilin proteins. GAPDH served as the loading control. **(C)** Bar graphs showing the relative fold change in viral and host proteins expression with respect to DMSO control. **(D)** Confocal microscopy images showing the levels of E2 and phalloidin during CHIKV infection.

In order to confirm the involvement of actin fibers in CHIKV progeny release, Vero cells were virus infected and drug treated as mentioned above and cells were fixed at 18 hpi. Thereafter, phalloidin staining was carried out to stain actin fibers in cells as it has been reported that fluorescent dye-labeled phalloidin stains only the actin fibers, but not the monomers (38). Phalloidin staining was found to be more prominent in infected cells without CMPD1 treatment and was more diffusely stained in CMPD1 treated infected cells. Furthermore, the expression pattern of CHIKV E2 protein was unchanged in both the samples as shown in Fig 4D. Taken together, the results depict that CHIKV utilizes the actin polymerization process for its progeny release through activation of MK2/MK3; however CMPD1 abrogates the whole process by inhibiting MK2/3 activation.

### CMPD1 inhibits CHIKV infection *in vivo*

In order to assess the bio-availability of a drug/inhibitor, computer models have been used as a valid alternative to experimental procedures for prediction of ADME (Absorption, Distribution, Metabolism and Excretion) parameters (39). The SwissADME Web tool (www.swissadme.ch) is one such tool which enables the computation of key physicochemical, pharmacokinetic, drug-like and related parameters for one or multiple molecules (40). Hence, the bioavailability of CMPD1 was predicted through the SwissADME tool and it was found that CMPD1 has high GI (Gastro Intestinal) absorption with a bioavailability score of 0.55 as shown in “S2 Table”.

In order to assess the antiviral effect of CMPD1 on CHIKV infection *in vivo*, 10-14 days old male C57BL/6 mice (n=5 per group) were infected with the IS strain and serum as well as tissue samples were harvested as per the protocol mentioned above. Viral RNA was isolated from the pooled serum samples (from respective group) and RT-PCR was carried out to amplify E1 gene of CHIKV. It was observed that the viral copy number was reduced remarkably (90%) in CMPD1 treated CHIKV infected mice in comparison to control (Fig 5A). Next, to compare the extent of tissue inflammation due to CHIKV infection in presence of drug, muscle tissue sections (from the site of injection) of the sacrificed mice at 4dpi were stained using Haematoxylene and Eosin and it was found that the infiltration of immune cells were less in CMPD1 treated tissue in comparison to control as shown in Fig 5B. Furthermore, to determine the relative levels of different cytokines/chemokines in CMPD1 treated mice, proteome profiling was carried out with the pooled serum samples as described above. It was noticed that the expressions of few selective inflammatory cytokines/chemokines, like CXCL13, RAGE, FGF and MMP9 were significantly reduced in CMPD1 treated mice sera, as shown in Fig 5C and 5D. Interestingly, HGF was upregulated in CMPD1 treated mice. To assess the protective action of CMPD1, survival curve analysis was performed. For that, CHIKV infected mice (5 per group) were fed with CMPD1 (5mg/kg) orally at 3hrs post CHIKV infection and then for 3 consecutive days at an interval of 24 hrs. The disease scoring was performed based on the symptoms described in the methods section and shown in “S3 Table”. Moreover, from the survival curve analysis as shown in Figure 5e, there was 100% mortality of the untreated CHIKV infected mice after 8 days post infection. In contrast, no mortality was observed for the CMPD1 treated CHIKV infected mice even after 8 days post infection. The data suggests that CMPD1 shows anti-CHIKV activity *in vivo*.

**Fig 5:**
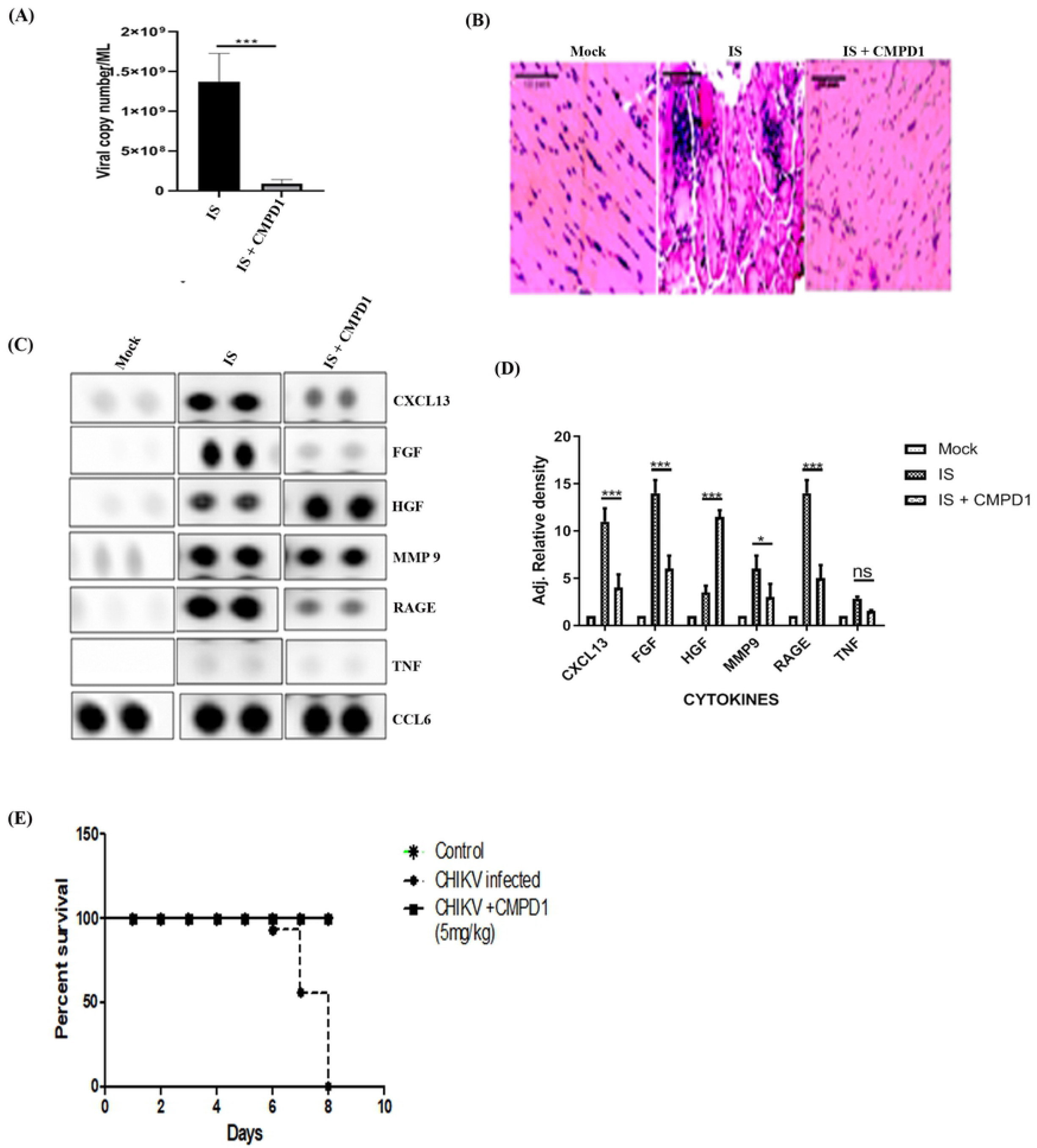
CMPD1 inhibits CHIKV infection in mice. **(A)** Bar graph showing the viral copy numbers in CHIKV infected and CMPD1 treated mouse serum samples. (B) H and E staining of mouse tissue samples with CHIKV infection and in presence/absence of CMPD1 **(C)** Array blot showing the expression of different cytokines after CHIKV infection in presence and absence of CMPD1. **(D)** Bar graph showing the relative band intensities of selected cytokines in mock, CHIKV infected and CMPD1 treated samples. **(E)** Survival curve showing the effect of CMPD1 in CHIKV infected mice

### CMPD1 modulates HSV and SARS CoV2 infection *in vitro*

In order to assess the efficacy of CMPD1 against other viruses like HSV-1 and SARS-CoV-2, Vero cells were infected with 0.1 MOI of HSV and SARS-CoV-2 separately and treated with different concentrations of CMPD1 post infection. The cells were incubated for 22 hpi and distinct morphological changes were visible under microscope between infected and drug treated cells as shown in Fig 6A and 6C. In case of HSV-1, the supernatants of infected and drug treated cells were harvested at 22 hpi and plaque assay was carried out to estimate the viral titers. It was observed that there was around 45% inhibition with 25μM and 90% inhibition in viral titers with 50μM of CMPD1 as shown in Fig 6B. However, in case of SARS-CoV-2, viral RNA was extracted from the supernatants at the same time point and cDNA synthesis was carried out. Then, the Nucleocapsid (NC) gene of SARS-CoV-2 was amplified by qRT-PCR using the gene specific primers [Forward: GTAACACAAGCTTTCGGCAG and Reverse:-GTGTGACTTCCATGCCAATG]. The copy number of the virus was calculated from the Ct value using the standard curve. It was observed that there was around 60% reduction with 50μM of CMPD1 and 88% reduction with 75 μM of CMPD1 as shown in Fig 6 D. The results indicate that CMPD1 is effective against other viruses like HSV-1 and SARS CoV2 *in vitro.*

**Fig 6:**
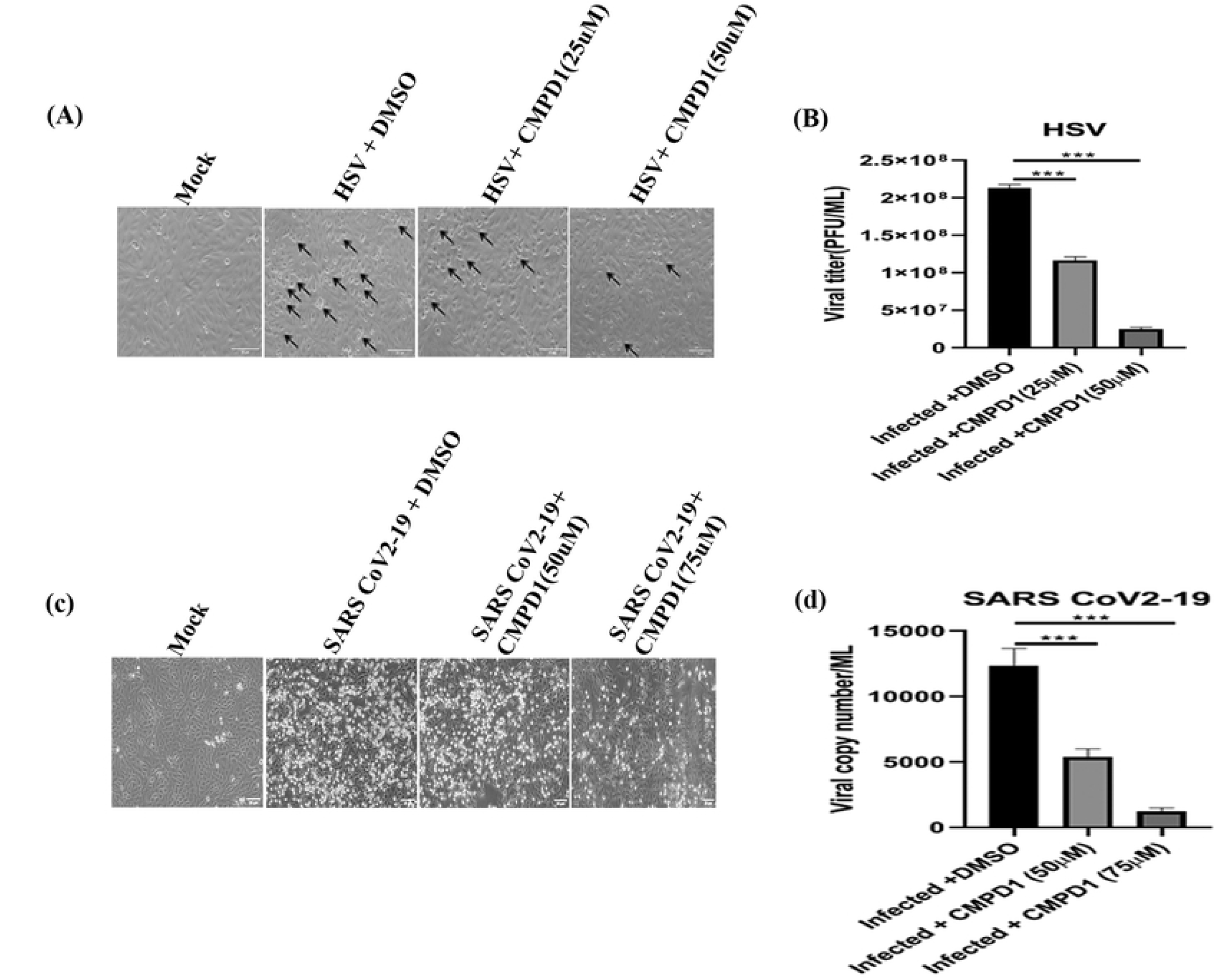
CMPD1 modulates HSV-1 and SARS-CoV-2 infection *in vitro*. Vero cells were infected with HSV-1 and SARS-CoV-2 (MOI 0.1) and treated with different concentrations of CMPD1. **(A and C)** Bright field images showing CPE in presence of CMPD1 for HSV-1 and SARS-CoV-2 infection. Black arrows indicate infected cells for HSV-1 and glowing cells represent SARS-CoV-2 infected cells. **(B and D)** Bar graph showing viral titer/copy number in presence of CMPD1 for HSV-1 and SARS-CoV-2 infection respectively.

## Discussion

CHIKV is now considered as a major public health concern. Due to lack of therapeutics and vaccine, a number of studies have been initiated to understand the function of viral proteins and the mechanisms of virus-mediated manipulation of host machineries for successful replication (41–43). The current investigation, aims to determine how host proteins are modulated during CHIKV infection in mammalian cell lines. In this regard, microarray analysis was carried out for mock and CHIKV-infected Vero cells and two genes MK2 and MK3 belonging to P38MAPK pathway were selected for further analysis.

Gene silencing of MK2 and MK3 abrogated around 58% CHIKV progeny release from the host cell and a MK2 activation inhibitor (CMPD1) treatment demonstrated 68% inhibition of viral infection suggesting a major role of MAPKAPKs during late CHIKV infection *in vitro*. Further, it was observed that the inhibition in viral infection is primarily due to the abrogation of lamellipodium formation through modulation of factors involved in the actin cytoskeleton remodeling pathway which is essential for virus release. Moreover, CHIKV-infected C57BL/6 mice demonstrated reduction in the viral copy number, lessened disease score and better survivability after CMPD1 treatment. In addition, reduction in expression of key pro-inflammatory mediators such as CXCL13, RAGE, FGF, MMP9 and increase in HGF (a CHIKV infection recovery marker) was observed indicating the effectiveness of this drug against CHIKV. Additionally, CMPD1 also inhibited the HSV1 and SARS CoV2-19 infection *in vitro*.

The roles of MK2 and MK3 have been implicated in few other viruses like Dengue (DENV), Murine Cytomegalovirus (MCMV), Kaposis Sarcoma Herpes Virus (KSHV), Rous Sarcoma Virus (RSV), Influenza A and Human Immunodeficiency Virus (HIV). In DENV, it was found that SB203580 (a P38MAPK inhibitor) treatment significantly reduced the phosphorylation of MAPKAPK2 and other substrates such as HSP27 and ATF2 which reduced DENV-induced liver injury in mice (44). In the case of MCMV, MK2 was reported to regulate cytokine responses towards acute infection, via IFNARI-mediated pathways and prevents formation of intrahepatic myeloid aggregates during infection (45). For KSHV, it was observed that the viral Kaposin B (KapB) protein binds and activates MK2, thereby selectively blocking decay of AU-rich mRNAs (ARE-mRNAs) that encode pro-inflammatory cytokines and angiogenic factors during latent KSHV infections (46). Furthermore, it was noticed that during RSV infection, pP38 is sequestered inside cytoplasmic inclusion bodies (IBs) resulting in substantial reduction in accumulation of MK2 and suppressing cellular responses to virus infection. Additionally, CMPD1 treatment reduced viral protein expression suggesting the importance of pMK2 in RSV protein translation (32). In case of Influenza A, it was observed that MK2 and MK3 are activated on virus infection enabling the virus to escape the antiviral action of PKR (47). Recently, it has been shown that CCR5-tropic HIV induces significant reprogramming of host CD4+ T cell protein production pathways and induces MK2 expression upon viral binding to the cell surface that are critical for HIV replication in host cells (48). However, reports pertaining to the involvement of MK2 and MK3 in alphavirus infection are not available. Hence, this investigation is one of the first to highlight the importance of MK2 and MK3 in CHIKV.

According to the results, it can be suggested that both MK2 and MK3 play important roles in CHIKV progeny release during CHIKV infection. After CHIKV infection, MK2 is phosphorylated which in turn phosphorylates LIMK1.(37) The LIMK1 then inactivates Cofilin by phosphorylating it (36) This results in accumulation of more p-Cofilin inside the cell than active Cofilin. As a result, Cofilin is unable to cleave the actin filaments into monomers. This leads to polymerization of actin filaments and subsequent lamellipodia formation which results in effective CHIKV progeny release, as shown in Fig 7A. However, CMPD1 treatment abrogates MK2/3 phosphorylation as a result of which LIMK is not able to inactivate Cofilin. Active Cofilin then cleaves actin polymers to monomers, thereby preventing lamellipodium formation and subsequent viral progeny release as shown in Fig 7B. Furthermore, *in vivo* studies demonstrate that CMPD1 treated mice do not develop complications post CHIKV infection. This can be speculated by the reduction in the expressions of certain virus induced inflammatory chemokines and cytokines like CXCL13, RAGE and FGF in CMPD1 treated mice sera. The involvements of these chemokines and cytokines have been reported for other virus infections before (49–51). Additionally, the expression of MMP9, a host factor involved in the degradation of extracellular matrix thereby promoting viral spread to neighbouring tissues **(51)** was also reduced in drug-treated samples indicating abrogation of viral transmission during CMPD1 treatment. In contrast, the expression of HGF (a known marker for CHIKV recovery during acute infection) (52) was upregulated during CMPD1 treatment thereby showing the effectiveness of CMPD1 against CHIKV in a mouse model. Nevertheless, it would be interesting to understand the detailed mechanism and role of these factors during CHIKV infection in future.

**Fig 7:**
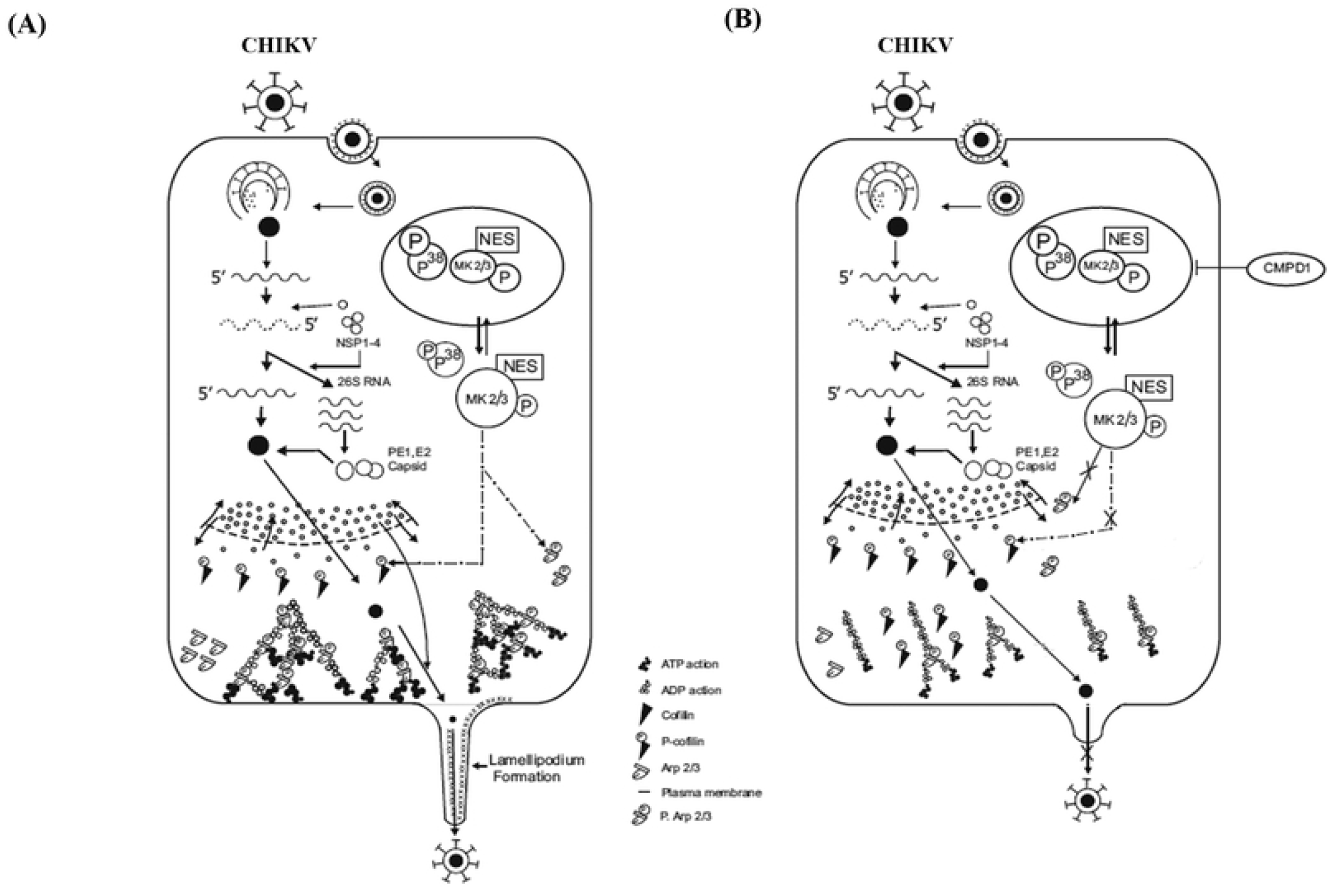
Proposed model for CHIKV infection. **(A)** During CHIKV infection, MK2/3 gets phosphorylated by P38 MAPK thereby exposing the Nuclear Export Signal (NES) of MK2. The phosphorylated forms of MK2/MK3 translocate to the cytoplasm and help in inactivating Cofilin through phosphorylation via LIMK-1 thereby promoting actin polymerization and lamellipodium formation. **(B)** Addition of CMPD1 blocks the phosphorylation of MK2, thereby blocking Cofilin phosphorylation and eventually inhibiting lamellipodium formation and CHIKV progeny release.

Thus, the current study highlights the importance of MK2 and MK3 (substrates of the p38MAPK pathway) as novel host factors involved during CHIKV infection. It also demonstrated CMPD1 as a novel inhibitor of CHIKV infection; hence, CMPD1 can be pursued as a potential lead for developing anti-CHIKV molecule to regulate disease manifestations.

## Supplementary information

S1 Table: - Differently modulated host genes for DRDE-06 classified into different metabolic pathways.

S2 Table: - Bioavailability prediction of CMPD1 through SWISSADME web tool.

S3 Table: - Disease scoring of CHIKV infected and drug treated mice

## Authors Contribution

SoC, SuC and PM conceived the idea and designed the experiments; PM, TKN, AK, SK, SC, AD, SD and SM performed wet lab experiments; SoC and SuC contributed reagents; SoC, SuC and PM analyzed and interpreted the data; SoC, PM, TKN, EL, AR and SuC wrote the manuscript. All authors read and approved the final version of this manuscript.

## Acknowledgement

We are thankful to Dr. M.M. Parida, DRDE; Gwalior, India for kindly providing IS and PS strains of CHIKV and Vero cell line. We like to acknowledge Dr. Roger Everett, Glasgow University, Scotland for providing the KOS strain of HSV-1. We are also grateful to Dr. Rupesh Dash of ILS, Bhubaneswar, India for providing the HEK293T cell line.

## Transparency Declarations

Nothing to declare.

